# Treatment Outcome of Diabetic Ketoacidosis Among Patients Atending General Hospital in North-West Ethiopia: Hospital Based Study

**DOI:** 10.1101/441964

**Authors:** Tadesse Melaku Abegaz, Gizework Alemnew Mekonnen, Eyob Alemayehu Gebreyohannes, Kasssahun Alemu Gelaye

## Abstract

**Background:** Diabetic ketoacidosis is an acute life-threatening complication of diabetes mellitus. There was limited data on level of in-hospital mortality, hospital stay and factors associated with length of hospital stay among diabetic patients admitted to diabetic ketoacidosis at Debretabor General Hospital.

**Objective:** The aim of the study was to determine the length of hospital stay and in-hospital mortality of diabetic ketoacidosis patients and to assess determinants of long hospital stay among diabetic patients admitted with Diabetic ketoacidosis at Debretabor General Hospital.

**Method:** A retrospective study was conducted at Debretabor General Hospital from June 1to 30, 2018. Participants included in the study were all diabetic patients with diabetic ketoacidosis admitted to the hospital from August 2010 to May 31, 2018 whose medical records contained complete pertinent data. The primary outcome was to determine the length of hospital stay and in-hospital mortality of diabetic ketoacidosis patients. All the statistical data was carried out using Statistical Package for Social Sciences (SPSS). Descriptive statistics was presented using means with standard deviation and percentages.

**Result:** A total of 387 patients’ medical records contained pertinent complete information included in this study. Mean age of the patients was 33.30± 14.96 years. The majority of patients were females 244 (63.0%). The mean length of hospital stay was 4.64(±2.802) days. About twenty percent 79(20.41%) patients had long hospital stay (>7days). The majority 370 (95.60%) of patients improved and discharged and 17 (4.40%) patients died in the hospital. patients who had mild DKA; AOR: 0.16 [0.03-0.78] and patients between the age of 35-44years, AOR: 0.125[0.017-0.92] had reduced length of hospital stay. further, patients with DKA precipitated by infection were 4.59 times more likely to have long hospital stay than patients with DKA precipitated by unknown causes; AOR 4.59[1.08-19.42].

**Conclusions:** In the current study, the mean length of hospital stay was around five days. About twenty percent patients had long hospital stay. Nearly ninety five percent of patients improved and discharged. The presence of infection, frequent rebound hyperglycemia and severity of DKA were the major determinants of long hospital stay.

## Background

Diabetes mellitus describes a group of metabolic disorders characterized by increased blood glucose concentration(1). People living with diabetes have a higher risk of morbidity and mortality than the general population. The international diabetes federation estimated that 415 million adults were diabetic in 2015(2). These patients are at risk of developing acute diabetic complications such as diabetic ketoacidosis (DKA). Diabetic ketoacidosis is an acute life threatening complication of diabetes mellitus (DM). Earlier thought to be a hallmark of type 1 diabetes mellitus (DM), DKA now is being recognized in type 2 DM also(3–6). In DKA, reduced effective insulin concentrations and increased concentrations of counter regulatory hormones lead to hyperglycemia and ketosis (7, 8). The classical clinical picture DKA includes a history of polyuria, polydipsia, weight loss, vomiting, dehydration, weakness, and mental status change along with pertinent physical findings (7, 9, 10).

DKA is a frequently encountered hospital emergency and is associated with significant morbidity and mortality. Its mortality outcome has been reported to be less than 5% in treatment experienced centers of the Americas, Europe and Asia (9). In Africa the mortality of DKA is unacceptably high with a reported death rate of 26 to 29% in studies from Kenya, Tanzania and Ghana (6). DKA is responsible for more than 500,000 hospital days per year at an estimated annual direct medical expense and indirect cost of 2.4 billion $ dollar. The triad of uncontrolled hyperglycemia, metabolic acidosis, and increased total body ketone concentration characterizes DKA (7, 11). In Ethiopia, mortality from hyperglycemic emergencies is not well documented but comparatively high in few studies. A study done in Jimma found a mortality rate of 9.8 % from hyperglycemic emergencies (12). Socioeconomic factors, particularly the cost and unreliability of insulin supplies are the major obstacles to the control of diabetes and prevention of ketoacidosis in Ethiopian patients(13). Mortality rate is lower in patients that received appropriate treatment and management of precipitating factors(14).

Pool of evidence highlighted that the prolonged hospital stay has been one of the major challenge in patients hospitalized with DKA. For instance, a hospital-based descriptive study done in Nepal in 2010 indicated that the average length of hospitalization was around 30 up to 60 days (15). Another hospital based observational study done in India showed that the duration of hospital stay ranged from 8- was 12 days (16). Data from Africa also suggested that patients stayed for more than ten days after admission with DKA (17) (18).

Length of hospital stay (LHS) and overall in hospital mortality (IHM) are the major primary outcomes that should be measured in DKA patients. Nonetheless, the data related to LHS and the rate of IHM due to DKA remain to be scarce in Ethiopia. A study highlighting LHS and overall IHM is required to improve management of DM and DKA related complications. Therefore, the current study aimed to determine the outcome of DKA and associated factors among diabetic patients admitted to Debretabor General Hospital (DGH), northwest Ethiopia.

## Methods

### Study setting, design and period

Hospital based cross-sectional study was done at DGH from 1 June to 30 June 2018. DGH is a government hospital located in northwest Ethiopia. The hospital diabetic clinic receives DM patients for follow up twice weekly as outpatients’ services. Accordingly, dose adjustment and regimen change is made routinely made based on the level of glucose control. DKA patients are also admitted for in patients care in the hospital.

### Population and

All DM patients with DKA who were admitted to inpatient ward of DGH were our source population. Particularly, patients with age ≥15 years old whose medical records contained complete pertinent data included. No forma sample size calculation was required as All DKA patients who were admitted to the inpatient ward of DGH and fulfilled the inclusion criteria August 2010 to May31 2018 were included.

### Study variables

Length of hospital stay and overall in hospital mortality were the independent variables. The Independent variables include: age, gender, residence, family history of DM, and class of DM, past admission due to DKA, severity of DKA, admission blood glucose readings, blood pressure, respiratory rate, pulse rate, co morbidities, precipitating factors, frequency of serum glucose rebound, and frequency of ketone rebound.

### Data quality control technique

Data collectors were trained intensively on contents of questionnaire, data collection methods and ethical concerns. The questionnaires was pre-tested on five percent of subjects and modified accordingly.

### Data collection methods

Medical record of patients with DKA admitted to the hospital was traced from patient logbook and drawn from card room. Selection of medical records for sampling was based on the physician’s confirmed diagnosis on patient logbooks. Participants included in the study were all diabetic patients with DKA admitted to DGH with age ≥15 years old whose medical records contained complete pertinent data. The data was collected by trained data collector using structured and pretested data extraction tool. Data was collected on patient demographics, presenting symptoms, precipitating causes of DKA, vital signs, biochemical profiles (admission blood glucose, admission urine ketone, urine glucose) at presentation to the inpatient department, time from presentation to resolution of urine ketones, length of hospitalization and treatment outcomes.

### Data Analysis

All the statistical data was conducted using Statistical Package for Social Sciences (SPSS), version 22 (SPSS Inc., Cary, NC, USA). Descriptive statistics was carried out for categorical variables. Binary logistic regression was done to determine the factors that affect length of hospital stay.

### Ethics

The study was conducted after ethical clearance letter received from research and ethics review committee of school of pharmacy, University of Gondar College of medicine and health science, hospital clinical director and head of the medical ward of DGH. Data was collected anonymously.

### Operational definitions

**DKA** is defined as admission blood glucose >250 mg/ dL and presence of ketonemia and/or ketoneuria (7).

**Hyperglycemia** is defined as random plasma glucose >200 mg/dL(19) and Hypoglycemia is defined as a blood glucose level ≤70 mg/dL(20).

**Euglycemia** is defined as serum glucose of between 100 and 200mg/dl(7).

**Normal** blood pressure is systolic of less than 120 and diastolic of less than 80 (120/80)(21)

**Serum glucose rebound** is defined as an increase in serum glucose level while the patient was admitted and on treatment of DKA.

**Long hospital stay** was defined as hospital stay for more than seven days and short hospital stay was considered if the patient stayed in the hospital for ≤ 7 days(22).

**Treatment outcomes** defined as the level of glycemic control, the length of hospital stay and in-hospital mortality of diabetic ketoacidosis.

**Poor glycemic control** is defined as a serum glucose rebound at least one times while the patient was on DKA treatment.

**Treatment outcome:** the length of hospital stay and in hospital mortality were the measures of treatment outcome in the context of the present study

### Results

#### Sociodemographic characteristics of DKA patients

About 387 patients were recruited in the study. Majority 305 (78.8%) of the patients had type 1 DM. around 244(63.0%) were females and 143(37.0%) were males. The mean age of the patients was 33.30 with standard deviation of 14.96, and it ranged from 15 to 64 years. More than two third of the participants 264 (68.2%) came from urban areas 123(31.8%). Nearly one-tenth of 50 (12.9%) patients reported facility history of DM

Patients lived with DM chronically for a mean duration of 26.21 (±39.62) months. The mean the frequency of DKA was found to be 1.5 times in a single patients with a maximum frequency of eight DKA episodes (table 1).

**Table 1:** Sociodemographic characteristics of DKA patients admitted at DGH from August 2010 to May 31, 2018.

| Characteristics | Frequency (%) |
| --- | --- |
| Age (mean ±SD) | 33.29 ($\pm 14.96$) |
| Sex |  |
| Male | 143(37.00) |
| Female | 244(63.00) |
| Residence |  |
| Urban | 264(68.20) |
| Rural | 123(31.80) |
| Family history |  |
| Yes | 50(12.90) |
| No | 53(13.70) |
| Unknown | 284(73.40) |
| Type of DM |  |
| Newly diagnosed type 1 dm | 146(37.70) |
| Newly diagnosed type 2 dm | 21(5.40) |
| Known type 1 dm | 159(41.10) |
| Known type 2 dm | 61(15.80) |
| Duration of DM(mean±SD) | 26.22($\pm 39.67$) |
| Mean frequency of DKA episode | 1.50($\pm 1.00$) |
| 1 times | 272(70.30) |
| 2 times | 74(19.10) |
| 3 times | 20(5.20) |
| $\geq 4$ times | 21(5.50) |

#### Length of hospital and in hospital mortality

The mean length of hospital stay was 4.64(±2.802) days. The minimum duration of stay was 1 day and the maximum was 18 days. About twenty percent 79(20.41%) patients had long hospital stay (>7days). The majority 370 (95.60%) of patients improved and discharged and 17 (4.40%) patients died in the hospital (Table 2). The rate of overall in-hospital mortality was found to be 17(4.5%) (Figure 1).

**Figure 1:**
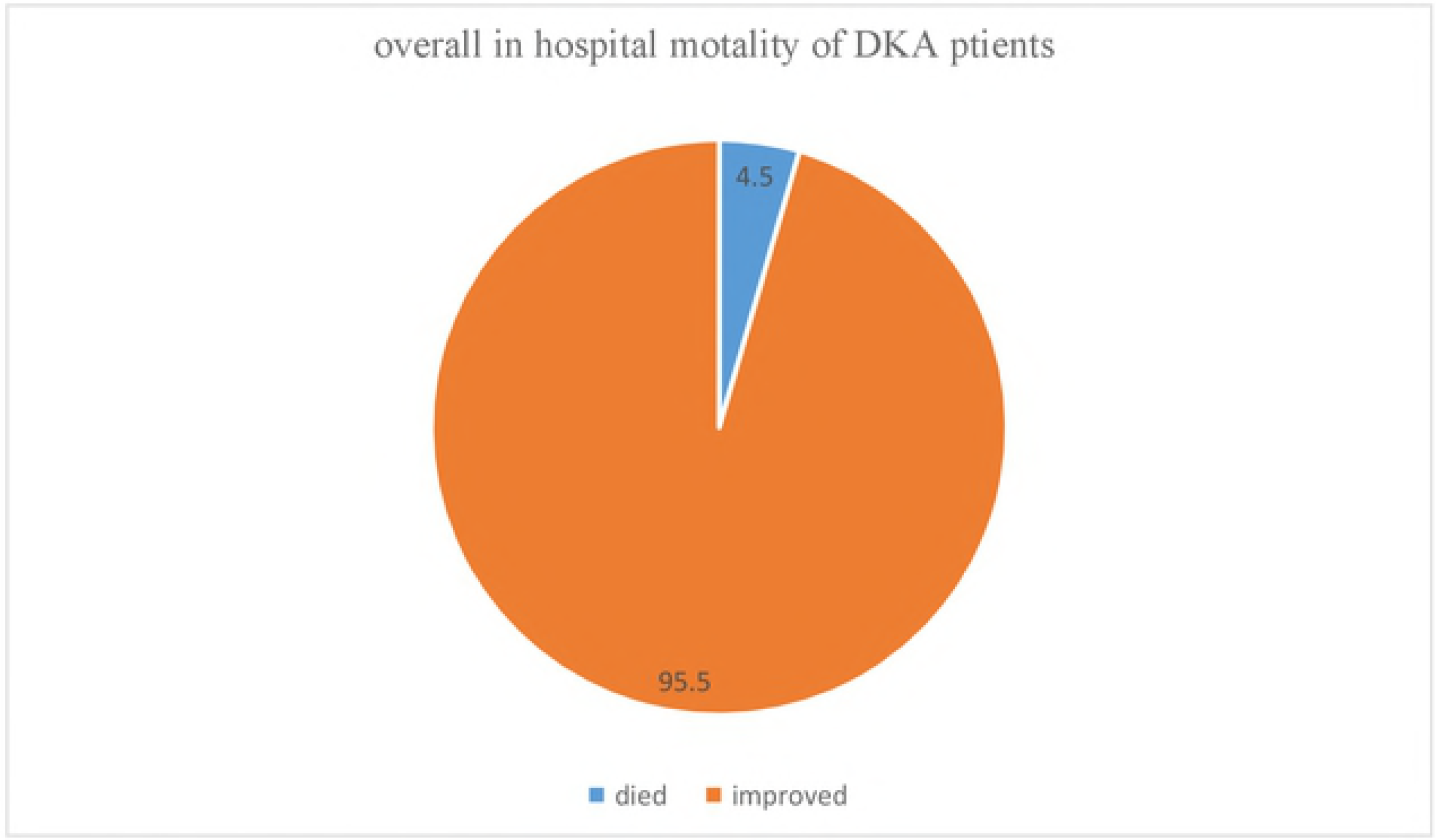
overall in-hospital mortality of DKA patients

**Table 2:** Length of hospital stay and treatment outcome in patients admitted at DGH from August
2010 to May 31, 2018.

| Length of hospital stay and treatment outcome | Frequency (%) |
| --- | --- |
| Mean length of hospital stay | 4.64(±2.80)days |
| Minimum length of hospital stay | 1 days |
| Maximum length of hospital stay | 18 days |

#### Determinants factors for length of hospital stay

Different factors have been studied as determinants of long hospital stay in DKA patients. It was found that patients who had mild DKA showed 84% reduction in the probability of long hospital stay than that of patients who had severe DKA; AOR: 0.16 [0.03-0.78] and patients with moderate DKA showed 83% less likelihood to have long hospital stay; AOR : 0.17[0.03-0.96]. Patients between the age of 35-44 had 87.5% chance of reduction on long hospital stay than patients of age 55-64 years; AOR: 0.125[0.017-0.92]. further, patients with DKA precipitated by infection were 4.59 times more likely to have long hospital stay than patients with DKA precipitated by unknown causes; AOR 4.59[1.08-19.42]. In addition, as the number of times the serum rebound glucose increase by one unit the likelihood of long hospital stay increased by more than 2.15 times; AOR: 2.15(1.76-2.63) (Table 3).

**Table 3:**
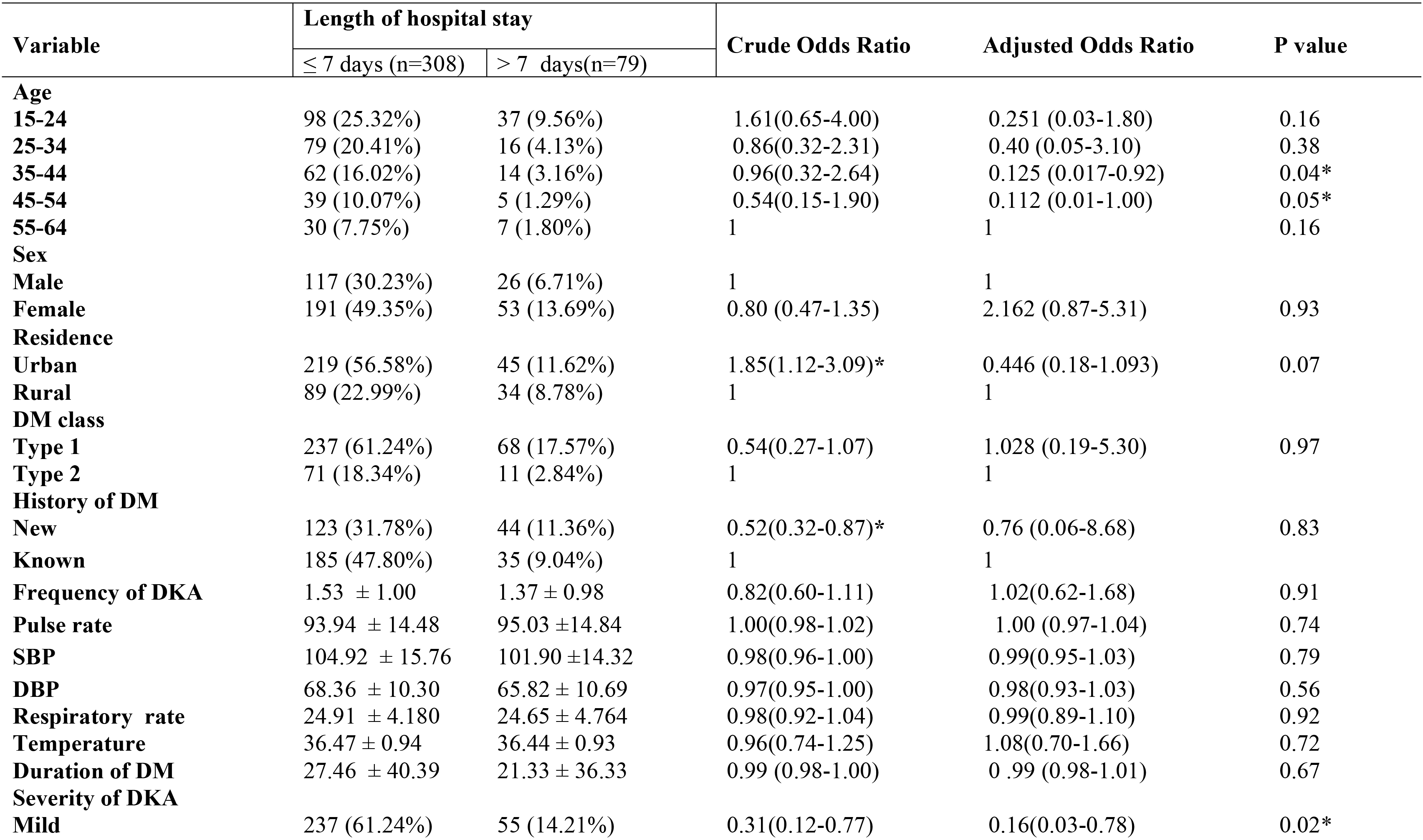
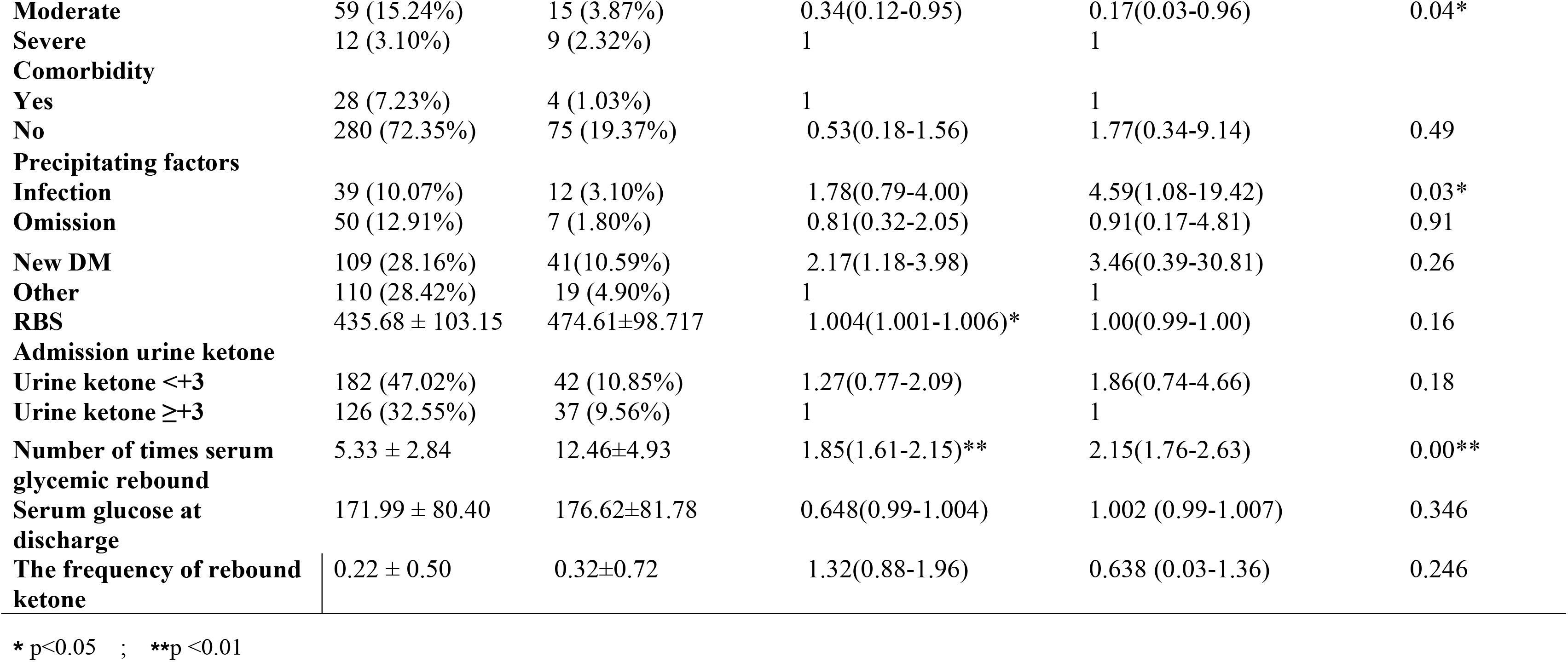
Determinants of long hospital stay in DKA patients at DGH from August 2010 to May 31, 2018.

| Variable | Length of hospital stay |  | Crude Odds Ratio | Adjusted Odds Ratio | P value |
| --- | --- | --- | --- | --- | --- |
|  | ≤ 7 days (n=308) | > 7 days(n=79) |  |  |  |
| **Age** |  |  |  |  |  |
| **15-24** | 98 (25.32%) | 37 (9.56%) | 1.61(0.65-4.00) | 0.251 (0.03-1.80) | 0.16 |
| **25-34** | 79 (20.41%) | 16 (4.13%) | 0.86(0.32-2.31) | 0.40 (0.05-3.10) | 0.38 |
| **35-44** | 62 (16.02%) | 14 (3.16%) | 0.96(0.32-2.64) | 0.125 (0.017-0.92) | 0.04* |
| **45-54** | 39 (10.07%) | 5 (1.29%) | 0.54(0.15-1.90) | 0.112 (0.01-1.00) | 0.05* |
| **55-64** | 30 (7.75%) | 7 (1.80%) | 1 | 1 | 0.16 |
| **Sex** |  |  |  |  |  |
| **Male** | 117 (30.23%) | 26 (6.71%) | 1 | 1 |  |
| **Female** | 191 (49.35%) | 53 (13.69%) | 0.80 (0.47-1.35) | 2.162 (0.87-5.31) | 0.93 |
| **Residence** |  |  |  |  |  |
| **Urban** | 219 (56.58%) | 45 (11.62%) | 1.85(1.12-3.09)* | 0.446 (0.18-1.093) | 0.07 |
| **Rural** | 89 (22.99%) | 34 (8.78%) | 1 | 1 |  |
| **DM class** |  |  |  |  |  |
| **Type 1** | 237 (61.24%) | 68 (17.57%) | 0.54(0.27-1.07) | 1.028 (0.19-5.30) | 0.97 |
| **Type 2** | 71 (18.34%) | 11 (2.84%) | 1 | 1 |  |
| **History of DM** |  |  |  |  |  |
| **New** | 123 (31.78%) | 44 (11.36%) | 0.52(0.32-0.87)* | 0.76 (0.06-8.68) | 0.83 |
| **Known** | 185 (47.80%) | 35 (9.04%) | 1 | 1 |  |
| **Frequency of DKA** | 1.53 ± 1.00 | 1.37 ± 0.98 | 0.82(0.60-1.11) | 1.02(0.62-1.68) | 0.91 |
| **Pulse rate** | 93.94 ± 14.48 | 95.03 ± 14.84 | 1.00(0.98-1.02) | 1.00 (0.97-1.04) | 0.74 |
| **SBP** | 104.92 ± 15.76 | 101.90 ± 14.32 | 0.98(0.96-1.00) | 0.99(0.95-1.03) | 0.79 |
| **DBP** | 68.36 ± 10.30 | 65.82 ± 10.69 | 0.97(0.95-1.00) | 0.98(0.93-1.03) | 0.56 |
| **Respiratory rate** | 24.91 ± 4.180 | 24.65 ± 4.764 | 0.98(0.92-1.04) | 0.99(0.89-1.10) | 0.92 |
| **Temperature** | 36.47 ± 0.94 | 36.44 ± 0.93 | 0.96(0.74-1.25) | 1.08(0.70-1.66) | 0.72 |
| **Duration of DM** | 27.46 ± 40.39 | 21.33 ± 36.33 | 0.99 (0.98-1.00) | 0.99 (0.98-1.01) | 0.67 |
| **Severity of DKA** |  |  |  |  |  |
| **Mild** | 237 (61.24%) | 55 (14.21%) | 0.31(0.12-0.77) | 0.16(0.03-0.78) | 0.02* |
| **Moderate** | 59 (15.24%) | 15 (3.87%) | 0.34(0.12-0.95) | 0.17(0.03-0.96) | 0.04* |
| **Severe** | 12 (3.10%) | 9 (2.32%) | 1 | 1 |  |
| **Comorbidity** |  |  |  |  |  |
| **Yes** | 28 (7.23%) | 4 (1.03%) | 1 | 1 |  |
| **No** | 280 (72.35%) | 75 (19.37%) | 0.53(0.18-1.56) | 1.77(0.34-9.14) | 0.49 |
| **Precipitating factors** |  |  |  |  |  |
| **Infection** | 39 (10.07%) | 12 (3.10%) | 1.78(0.79-4.00) | 4.59(1.08-19.42) | 0.03* |
| **Omission** | 50 (12.91%) | 7 (1.80%) | 0.81(0.32-2.05) | 0.91(0.17-4.81) | 0.91 |
| **New DM** | 109 (28.16%) | 41(10.59%) | 2.17(1.18-3.98) | 3.46(0.39-30.81) | 0.26 |
| **Other** | 110 (28.42%) | 19 (4.90%) | 1 | 1 |  |
| **RBS** | 435.68 ± 103.15 | 474.61±98.717 | 1.004(1.001-1.006)* | 1.00(0.99-1.00) | 0.16 |
| **Admission urine ketone** |  |  |  |  |  |
| **Urine ketone <+3** | 182 (47.02%) | 42 (10.85%) | 1.27(0.77-2.09) | 1.86(0.74-4.66) | 0.18 |
| **Urine ketone ≥+3** | 126 (32.55%) | 37 (9.56%) | 1 | 1 |  |
| **Number of times serum**<br>**glycemic rebound** | 5.33 ± 2.84 | 12.46±4.93 | 1.85(1.61-2.15)** | 2.15(1.76-2.63) | 0.00** |
| **Serum glucose at**<br>**discharge** | 171.99 ± 80.40 | 176.62±81.78 | 0.648(0.99-1.004) | 1.002 (0.99-1.007) | 0.346 |
| **The frequency of rebound**<br>**ketone** | 0.22 ± 0.50 | 0.32±0.72 | 1.32(0.88-1.96) | 0.638 (0.03-1.36) | 0.246 |

\* p<0.05 ; \*\*p <0.01

When patients are followed for through their hospital stay, males tend to die more earlier than females did (log rank,:1.5, p-value: 0.04) (figure 2).

#### Discussions

This study aimed at describing the treatment outcome of patients with DKA who were admitted to the inpatient ward of DGH. Successful treatment of DKA requires a prompt correction of hyperglycemia, dehydration and electrolyte disturbance (7, 10, 11). However, patients usually tend to stay for reasonably long time in the hospital till these parameters are up to the normal level. if not, some patients die of complications experienced in the hospital. Our study revealed that the mean length of hospital stay to be approximately 5 days. In contrast, the average length of hospital stay was very high in research done in sub Himalayan region which was around 9 days (16), seven days in Saudi Arabia (23), nine days in Pakistan(24) and south Africa (25). However, our study found a relatively long hospital stay when compared with a research done in Australia and Newzeland (26). This discrepancy might be the due to the better management of the precipitating factors, adequate management of complications, better control of rebound glucose in the case of developed countries. Again, the coincidence of infections as precipitant might have contributed to prolonged hospital stay in our study. Our study showed that patients whose DKA precipitated by infection had long hospital stay than those precipitated by undefined precipitants this result was in line with a research done at Indian (27) The coincidence of infections and DM has multidirectional impact on the outcome of patients. Infectious diseases tended to be more frequent and serious in patients with diabetes DM, which automatically leads to morbio-mortality. High burden of infections in these patients is due to the hyperglycemic environment which confers immune dysfunction, decrease in the antibacterial activity of urine, and greater number of admissions. Often times, the length of hospital stay increases and the lives of patients deteriorates when they presented with infections (28) (29)

Many factors found to influence the length of hospital stay in DKA patients in the present study. It was indicated that the length of hospital stay was affected severity of DKA. Patients with mild DKA had shorter length of stay than patients with severe DKA. A retrospective cohort study done at Israel (30)showed that length of hospital stay were worse in the severe group. Also, a cross sectional observational study done in Bangladesh showed that patients with mild DKA took short duration to recover than severe DKA (31). Another cross-sectional study done in Libya (32) showed that Patients with severe episodes were admitted for slightly longer periods than non-severe cases. Patients with severe DKA are frequently presented with in severe acidotic state and severe alteration in the level of consciousness (3, 7) this might cause complication (33)and correcting severe DKA requires long period of time this might be the reason for long hospital stay in severe DKA patients.in this study revealed that frequent rebound of hyperglycemia was associated with prolonged hospital stay. The main stay therapy of DKA relies in the correction of hyperglycemia with the administration of insulin. However, the initiation and continuation of adequate amount of insulin depends on the level of expertise that the professional bears which might affect the rapid identification of DKA(34).

In the present study approximately five percent of patients died in the hospital which is almost equal to a research done in Israel 4.1 %(35). Mortality was very high 16.66% in a research conducted India (16), 7.5% seven in Zambia(36) and nine percent in another study done in India (37). Still the rate of mortality was relatively high in our study. The reason might be high prevalence of infection which was approximately 13.2% and treatment complications including hypoglycemia as well as comorbidities.

#### Strength and limitation of the study

In general, the current study was the first in its kind that assessed death rate and length of hospital stay of DKA patients admitted in DGH. The findings of the study will help clinician to improve the management of DKA in general Hospital. The study also included all eligible DKA patients. However, this study also had few limitations. Firstly, it was a retrospective review of records performed at one hospital that is difficult to generalize. In addition, there was no consistency in completion of clinical notes that may have contributed to an information bias.

### Conclusion

In the current study, the mean length of hospital stay was around five days. About twenty percent patients had long hospital stay. Nearly ninety five percent of patients improved and discharged. The presence of infection, frequent rebound hyperglycemia and severity of DKA were the major determinants of long hospital stay. Since, infection was highlighted as a determinant factor for long hospital stay, infection preventions might attenuate the frequent incidence of DKA and its untoward impact on the lives of DM patients.

## Abbreviation

AOR: Adjusted Odds Ratio
COR: Crude Odds Ratio
DGH: Debretabor General Hospital
DM: Diabetes Mellitus
IHM: In hospital Mortality
LHS: Length Hospital Stay
SD: Standard Deviation
SPSS: Statistical Package for Social Sciences
US: United States

## Acknowledgements

We would like to acknowledge university of Gondar for the overall support.

## Availability of data and materials

All the necessary data was include in the manuscript

## Ethics approval and consent to participate

Ethical clearance was secured from university of Gondar. Consent to participate was not required as the data was collected retrospectively.

## Consent for publication

Not applicable

## References

1. Association AD. Diagnosis and classification of diabetes mellitus. Diabetes care. 2014;37(Supplement 1):S81–S90.

2. Ogurtsova K, da Rocha Fernandes J, Huang Y, Linnenkamp U, Guariguata L, Cho N, et al. IDF Diabetes Atlas: Global estimates for the prevalence of diabetes for 2015 and 2040. Diabetes research and clinical practice.128:40–50.

3. Ibrahim AM. An Overview of Diabetic Ketoacidosis. EC Endocrinology and Metabolic Research. 2017;2(1):24–30.

4. Ranjan A, Thakur S, Mokta J, Bhawani R, M. G. Clinical profile of diabetic ketoacidosis in adults in sub himalayan region: hospital based study. International Journal of Basic and Applied Medical Sciences 2016;6(2):64–70.

5. Sushma Trikha, Neelima Singh, Uttarwar P. Triggers in Diabetic Ketoacidosis and Predictors of Adverse Outcome. IJAR. 2015;1(10):230–4.

6. Van Zyl DG. Diagnosis and treatment of diabetic ketoacidosis: CPD. South African Family Practice. 2008;50(1):35–9.

7. Kitabchi AE, Umpierrez GE, Miles JM, Fisher JN. Hyperglycemic crises in adult patients with diabetes. Diabetes care. 2009;32(7):1335–43.

8. Chiasson J-L, Aris-Jilwan N, Bélanger R, Bertrand S, Beauregard H, Ékoé J-M, et al. Diagnosis and treatment of diabetic ketoacidosis and the hyperglycemic hyperosmolar state. Canadian Medical Association Journal. 2003;168(7):859–66.

9. Association AD. Hyperglycemic crises in diabetes. Diabetes Care. 2004;27(suppl 1):s94–s102.

10. Goguen J, Gilbert J. Hyperglycemic emergencies in adults. Canadian journal of diabetes.37:S72–S6.

11. Yehia BR, Epps KC, Golden SH. Diagnosis and management of diabetic ketoacidosis in adults. Hospital Physician. 2008;35:21–6.

12. Desse TA, Eshetie TC, Gudina EK. Predictors and treatment outcome of hyperglycemic emergencies at Jimma University Specialized Hospital, southwest Ethiopia. BMC research notes.8(1):553.

13. Lester FT. Ketoacidosis in Ethiopian diabetics. Diabetologia. 1980;18(5):375–7.

14. Qari F. Clinical characteristics of patients with diabetic ketoacidosis at the Intensive Care Unit of a University Hospital. Pakistan journal of medical sciences. 31(6):1463.

15. Maskey R, Shakya DR, Nikesh B, Krishna KA, Lavaju P, Kattel V, et al. Clinical profile of diabetic ketoacidosis in tertiary care hospital of Eastern Nepal. Indian journal of endocrinology and metabolism.19(5):673.

16. A. Ranjan S, Thakur J, Mokta R, Bhawani, Garg M. Clinical Profile of Diabetic Ketoacidosis in Adults in SubHimalayan Region: Hospital Based Study. International Journal of Basic and Applied Medical Sciences ISSN: 2277-2103 2016 Vol. 6 (2) May-August, pp.;6(2):64.

17. Elmehdawi R, Elmagerhei H. Profile of diabetic ketoacidosis at a teaching hospital in Benghazi, Libyan Arab Jamahiriya. Eastern Mediterranean Health Journal 2010; 16(3).

18. Gninkoun CJ, Alassani AbSC, Sagna Y, Adjagba P, Djrolo Fo. Diabetic Ketosis Decompensations at the National Hospital in Benin (West Africa), What Did We Learn about the Precipitating Factors? Journal of Diabetes Mellitus.6(04):301.

19. Moghissi ES, Korytkowski MT, DiNardo M, Einhorn D, Hellman R, Hirsch IB, et al. American Association of Clinical Endocrinologists and American Diabetes Association consensus statement on inpatient glycemic control. Diabetes care. 2009;32(6):1119–31.

20. Association AD. Glycemic targets: standards of medical care in diabetes—2018. Diabetes Care. 2018;41(Supplement 1):S55–S64.

21. Whelton PK, Carey RM, Aronow WS, Casey DE, Collins KJ, Himmelfarb CD, et al. 2017 ACC/AHA/AAPA/ABC/ACPM/AGS/APhA/ASH/ASPC/NMA/PCNA guideline for the prevention, detection, evaluation, and management of high blood pressure in adults: a report of the American College of Cardiology/American Heart Association Task Force on Clinical Practice Guidelines. Journal of the American College of Cardiology. 2018;71(19):e127–e248.

22. Desse T, Eshetie T. Determinants of Long Hospital Stay among Diabetic Patients Admitted with Diabetic Ketoacidosis at Jimma University Specialized Hospital. J Trauma Stress Disor Treat 6.1:2.

23. Almalki MH, Buhary BM, Khan SA, Almaghamsi A, Alshahrani F. Clinical and Biochemical Characteristics of Diabetes Ketoacidosis in a Tertiary Hospital in Riyadh. Clinical Medicine Insights: Endocrinology and Diabetes.9:CMED. S39639.

24. Jabbar A, Farooqui K, Habib A, Islam N, Haque N, Akhter J. Clinical characteristics and outcomes of diabetic ketoacidosis in Pakistani adults with Type 2 diabetes mellitus. Diabetic medicine. 2004;21(8):920–3.

25. Ndebele NF, Naidoo M. The management of diabetic ketoacidosis at a rural regional hospital in KwaZulu-Natal. African journal of primary health care & family medicine. 2018;10(1):1–6.

26. Venkatesh B, Pilcher D, Prins J, Bellomo R, Morgan TJ, Bailey M. Incidence and outcome of adults with diabetic ketoacidosis admitted to ICUs in Australia and New Zealand. Critical Care. 2015;19(1):451.

27. Varadarajan P, suresh S. Role of Infections in Children with Diabetic Ketoacidosis- A Study from south india Int J Diabetes Clin Res 2014; 1:2.

28. Ko M-C, Chiu AW-H, Liu C-C, Liu C-K, Woung L-C, Yu L-K, et al. Effect of diabetes on mortality and length of hospital stay in patients with renal or perinephric abscess. Clinics. 2013;68(8):1109–14.

29. Casqueiro J, Casqueiro J, Alves C. Infections in patients with diabetes mellitus: A review of pathogenesis. Indian journal of endocrinology and metabolism. 2012;16(Suppl1):S27.

30. Barski L, Nevzorov R, Rabaev E, Jotkowitz A, Harman-Boehm I, Zektser M, et al. Diabetic ketoacidosis: clinical characteristics, precipitating factors and outcomes of care. IMAJ-Israel Medical Association Journal.14(5):299.

31. Nazneen S, Ahmed F, Ashrafuzzaman S, Uddin KN, Ahsan AA, Faruq MO, et al. Clinical Presentation and Biochemical Abnormalities in Patients Presented with Diabetic Ketoacidosis in BIRDEM Hospital. Bangladesh Critical Care Journal. 2017;5(1):7–10.

32. Elmehdawi RR, Ehmida M, Elmagrehi H, Alaysh A. Incidence and mortality of diabetic ketoacidosis in Benghazi-Libya in 2007. Oman medical journal.28(3):178.

33. Adnet F, Baud F. Relation between Glasgow Coma Scale and aspiration pneumonia. The Lancet. 1996;348(9020):123–4.

34. Jayashree M, Sasidharan R, Singhi S, Nallasamy K, Baalaaji M. Root cause analysis of diabetic ketoacidosis admissions at a tertiary referral pediatric emergency department in North India. Indian journal of endocrinology and metabolism. 2017;21(5):710.

35. Barski L, Nevzorov R, Rabaev E, Jotkowitz A, Harman-Boehm I, Zektser M, et al. Diabetic ketoacidosis: clinical characteristics, precipitating factors and outcomes of care. The Israel Medical Association journal: IMAJ.14(5):299–303.

36. Kakusa M, Kamanga B, Ngalamika O, Nyirenda S. Comatose and noncomatose adult diabetic ketoacidosis patients at the University Teaching Hospital, Zambia: Clinical profiles, risk factors, and mortality outcomes. Indian journal of endocrinology and metabolism.20(2):199.

37. Mahesh M, ShIvaSwaMy RP, ChandRa BS, Syed S. The Study of Different Clinical Pattern of Diabetic Ketoacidosis and Common Precipitating Events and Independent Mortality Factors. Journal of clinical and diagnostic research: JCDR.11(4):OC42.

